# Biological Aging of the Cardiopulmonary System

**DOI:** 10.64898/2026.06.09.731192

**Authors:** Dongnan Liu, Pramath Doddaballapur, Zhongyu Cai, Rira Choi, Jack Di Palo, Nicole Guerrera, Nebal S. Abu Hussein, Meredith S. Gwin, Liqin Lin, Siming Zheng, Yuening Zhang, Aurelien Justet, Abhay B. Ramachandra, Xiting Yan, Edward P. Manning

**Affiliations:** Department of Biostatistics, Yale School of Public Health, New Haven, Connecticut, USA; University of California Santa Barbara, Santa Barbara, California, USA; Department of Biomedical Engineering, Yale School of Medicine, New Haven, Connecticut, USA; Section of Pulmonary, Critical Care, and Sleep Medicine, Yale School of Medicine, New Haven, Connecticut, USA; Translational Research Imaging Center, Yale School of Medicine, New Haven, Connecticut, USA; Section of Pulmonary, Critical Care, and Sleep Medicine, Yale School of Medicine, New Haven, Connecticut, USA; Department of Respiratory and Critical Care Medicine, The First Affiliated Hospital of Zhengzhou University, Zhengzhou, Henan, China; Section of Pulmonary, Critical Care, and Sleep Medicine, Yale School of Medicine, New Haven, Connecticut, USA; Service de Pneumologie, Centre de Compétence Maladies Pulmonaires Rares–UMR 6030 CNRS-ISTCT, Université de Normandie, Caen, France; Department of Mechanical Engineering, Iowa State University, Ames, Iowa, USA; Section of Pulmonary, Critical Care, and Sleep Medicine, Yale School of Medicine, New Haven, Connecticut, USA; Department of Biostatistics, Yale School of Public Health, New Haven, Connecticut, USA; VA Connecticut Healthcare System, West Haven, Connecticut, USA; Section of Pulmonary, Critical Care, and Sleep Medicine, Yale School of Medicine, New Haven, Connecticut, USA

## Abstract

Age-related stiffening of large arteries is a predictor of cardiovascular morbidity and mortality, yet how pulmonary vascular stiffening integrates with right ventricular (RV) and lung functional decline—and how best to quantify “biological” cardiopulmonary aging—remains unclear. Here we map cardiopulmonary aging across the adult murine lifespan by integrating RV, proximal pulmonary artery (PA), and lung biomechanics with single-cell transcriptomics. Using *ex vivo* biaxial testing of the proximal PA, *in vivo* echocardiography, and lung mechanics, we find that cardiopulmonary aging is phase-dependent: PA circumferential stiffening and reduced distensibility progress largely linearly with age; whereas, RV remodeling and lung mechanical changes exhibit non-linear trajectories. This is consistent with early intrinsic functional decline of cells and organs followed by later, extrinsic load-dependent structural adaptation. To quantify organ-level biological aging, we apply principal component analysis to PA, RV, and lung feature sets to derive physiology-based aging scores that summarize coordinated variance within and across organs. Anchoring differential gene expression in PA single-cell RNA-seq to these continuous biological aging scores rather than chronological age reveals extensive, cell-type–specific remodeling programs (13,636 genes) that are sparse or non-informative when modeled by chronologic age. Biological aging associates across endothelia, smooth muscle cells, fibroblasts, and perivascular macrophages with increased oxidative phosphorylation signatures alongside suppression of adaptive/regulatory pathways, including impaired endothelial mechanotransduction, reduced smooth muscle Wnt signaling, altered extracellular matrix remodeling programs, and erosion of macrophage innate immune and TGFβ/NF-κB signaling nodes. These findings support a model in which pulmonary arterial stiffening is not merely a marker but an active contributor to cardiopulmonary aging via a biomechanical–metabolic–inflammatory uncoupling that diminishes vasoactive and mechano-adaptive reserve and promotes a positive feedback loop. Together, our work establishes physiology-derived biological aging as a powerful framework for interpreting vascular single-cell aging trajectories and identifies mechanistic pathways to target pulmonary vascular stiffening and preserve cardiopulmonary function with age.

## Introduction

There is an increasing appreciation of the negative physiological and clinical effects of vascular stiffness—the reduced ability of large arteries (e.g., the aorta) to expand and recoil with each heartbeat. Aortic stiffness increases systolic blood pressure and pulse pressure, raises the workload and oxygen demand of the left ventricle, and accelerates damage to delicate microvascular beds in organs such as the brain and kidneys; it also impairs the normal “cushioning” of pulsatile flow, promoting hypertension, left ventricular hypertrophy, heart failure, stroke, and chronic kidney disease^1-5^. While arterial stiffening tends to rise with chronological age, cardiovascular risk often tracks more closely with biological aging—the cumulative impact of genetics, lifestyle, inflammation, metabolic health (e.g., diabetes), smoking, sleep, and environmental exposures—because these factors can make vessels “older” or “younger” than a person’s calendar age^6^. In practice, two people of the same chronological age can have very different vascular stiffness and therefore very different risk, which is why measures that reflect biological age (including arterial stiffness) can be more informative for prevention and treatment than age in years alone.

In prior work, we demonstrated that the proximal pulmonary artery (PA) undergoes age-associated biomechanical remodeling characterized by a diminished capacity to store elastic energy and increased circumferential stiffness.^7^ These vascular changes were accompanied by reduced exercise capacity and declining function of both the lung and the right ventricle (RV), supporting the view that pulmonary vascular aging is tightly coupled to integrated cardiopulmonary performance.

At the molecular level, transcriptional profiling of the PA revealed aging-associated programs consistent with senescence across multiple vascular and perivascular cell populations, including perivascular macrophages, adventitial fibroblasts, and medial smooth muscle cells. Older pulmonary arteries exhibited increased expression of genes linked to extracellular matrix (ECM) turnover—encompassing components of the TGFβ signaling axis—and evidence of enhanced intercellular communication among macrophages, fibroblasts, and smooth muscle cells^7^. Collectively, these data suggested candidate biomarkers of pulmonary vascular aging and highlighted pathways that may be leveraged as mechanistic targets for anti-aging interventions.

However, the interpretability and translational reach of these findings were constrained by study design, most notably by modeling aging as a categorical rather than continuous exposure. We therefore posit that cardiopulmonary aging is intrinsically non-linear across the lifespan, from young adulthood through advanced age. Key structural, functional, and transcriptional transitions may occur in phases rather than as uniform trajectories. We further hypothesize that the PA is not merely a passive marker but an active driver of age-related cardiopulmonary decline, participating in a feedback loop in which PA remodeling alters RV afterload and pulmonary hemodynamics, thereby influencing RV and lung function, while concurrent changes in RV performance and lung mechanics reciprocally shape PA remodeling.

## Methods

### Mice

This study received approval from the Yale University Institutional Animal Care and Use Committee. Both female and male C57BL/6J WT mice (young and old) were acquired from Jackson Laboratory (Bar Harbor, ME), and additional old mice were sourced from the NIA Aging Mice Colony. The mice were housed in an antigen-free and virus-free animal care facility with a 12-hour light/dark cycle. It is recognized that mice of similar strains from different environments may exhibit phenotypic differences^8,9^. To minimize such environmental variations, some mice from Jackson Laboratory were inbred and aged locally. Mice from the NIA Aging Colony were procured at 20 months of age and further aged locally. The mice were provided with standard rodent chow and had free access to water. For ex vivo testing and transcriptomic analyses, the animals were euthanized with an overdose of urethane administered via intraperitoneal injection, followed by exsanguination and the harvesting of hearts, lungs, and pulmonary arteries.

### Voluntary Exercise

We measured the daily running distance of mice using in-cage running wheels (Actimetrics Wireless Low-profile Running Wheel Model ACE-557-WLP) with ClockLab Data Collection Software from Lafayette Instrument. These devices were designed to avoid interfering with normal housing conditions. Mice were given two days to acclimate to their cages equipped with the running wheel, and on the third day, we measured the distance run in meters over a 24-hour period. All mice participated in this measurement.

### Biomechanical Measurement and Analysis

Specimens were excised from the main pulmonary artery to the first branch of the right (RPA) and left (LPA) pulmonary artery, and prepared as previously described^10^. After flushing out the blood with a Hanks buffered saline solution^11^, perivascular tissue and fat were gently removed, and the LPA, ligamentum arteriosum, and small branch vessels were ligated with sutures. The RPA was cannulated onto custom glass micropipettes and secured with ligatures beyond the main pulmonary artery bifurcation and the first branch of the pulmonary artery at the other end. The specimen was submerged in Hanks buffered saline solution at room temperature to eliminate smooth muscle contractility and was biaxially tested using a custom computer-controlled testing device^12^.

To ensure reproducibility and rigor, we employed the same seven passive testing protocols as previously described^7,13^. These included pressure-distension tests at three fixed axial lengths (1.05, 1.00, and 0.95 times the in vivo length) and axial force-extension tests at four constant pressures (5, 15, 25, and 40 mmHg). Diameter was measured using a videoscope, length was controlled with a micro-stepper motor, and pressure and force were measured with standard transducers.

We modeled the passive mechanical behavior using a 2-D formulation, where the residual stresses homogenize the stress field, making mean values good estimates of overall wall stress^14^. The wall was modeled using a hyperelastic constitutive formulation consisting of a neo-Hookean term to capture elastin-dominated contributions and four-fiber families with Fung-like exponential behaviors to capture collagen and smooth muscle contributions. This model allows us to derive clinically and mechanobiologically relevant quantities such as biaxial wall stress and stiffness at different loads or deformations of interest. Details of the parameter estimation for these passive constitutive functions can be found in prior publications^10,13^.

Methods for measuring contractility in a biaxial setup are detailed in our previous work^10,15^. In this study, we report contractility at a constant distending pressure of 15 mmHg and subject-specific in vivo stretch under vasoactive stimulation with 100 mM potassium chloride (KCl) and 100 mM phenylephrine (PE), within an oxygenated Krebs physiological solution maintained at 37°C and pH 7.4.

Pulse wave velocity (PWV) serves as an integrated measure of an artery’s structural stiffness, which is influenced by both its geometry and material properties. PWV can be accurately approximated based on vessel distensibility (D), which is defined as

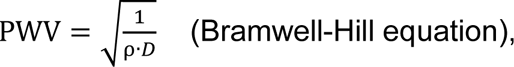

where ρ represents the mass density of the contained fluid (approximately 1050 kg·m^-3^) and D (in Pa^-1^or kg^-1^m·s^2^) is defined as the normalized change in arterial inner diameter from end-systole to end-diastole, divided by the change in end-systolic and end-diastolic pressures^16^. Pulmonary artery pulse wave velocity (PWV) and its impact on right ventricular (RV) hemodynamics can be assessed non-invasively using these variables^17-19^.

### Lung Mechanics Measurement and Analysis

Mice were anesthetized using urethane (1 g/kg administered in 10% solution with sterile water) and tracheostomized, then connected to the Flexivent system (FlexiVent®, SCIREQ©, Montreal, QC, Canada). Succinylcholine (1 mg/kg) was administered via intraperitoneal injection to eliminate spontaneous breathing. FlexiVent perturbations and oscillations were performed and analyzed using the FlexiWare Version 7.6 software, Service Pack 6 to obtain lung pressure-volume loops, static lung compliance (Cst), inspiratory capacity (IC), dynamic compliance (Crs), elastance (Ers), airway resistance (Rn), tissue dampening (G), parenchymal stiffness (H) and hysteresis. Maneuvers and perturbations continued until acquiring three suitable measurements. A coefficient of determination of 0.95 was the lower limit for suitable measurements. An average of three measurements for each metric was calculated per mouse.

### Ultrasonography

Noninvasive assessment of cardiac function was conducted in additional mice using transthoracic echocardiograms under light anesthesia (1.5% isoflurane) while maintaining physiological temperature^20^. Standardized cardiac views were obtained using a high-resolution ultrasound system (Vevo 2100, VisualSonics, Toronto, ON, Canada) equipped with an ultrahigh frequency (40 MHz) linear array transducer. B-mode two-dimensional (2D) images of the right ventricle (RV) and right atrium (RA) were captured from an apical four-chamber view, while images of the pulmonary artery were obtained from a parasternal short-axis view at the level of the aortic valve. Additionally, M-mode and tissue Doppler imaging (TDI) of the lateral tricuspid annulus were recorded from the apical four-chamber view. The pulmonic valve was visualized at the level of the leaflet tips using pulsed wave Doppler. Offline measurements included RV outflow tract (RVOT) diameter, tricuspid annular plane systolic excursion (TAPSE), RV systolic myocardial velocity (s’), pulmonary artery acceleration time (PAT), pulmonary ejection time (PET), two-dimensional end systolic (ES) and diastolic (ED) right ventricle area, and RA area. These measurements were performed using Vevo Lab software (version 3.2.6, VisualSonics) by an experienced sonographer. The fractional area change of the right ventricle was calculated as (ED-ES)/ED × 100^21^.

### Physiologic Measurement Modeling

Physiological data was plotted as a function of time (age in months). When we observed monotonic or curvilinear trends we modeled the trend using a best-fit curve in the form *y* = *y*_o_*e*^*kx*^. When rapid increases or decreases in data were observed preceded or followed by plateaus, we modeled that behavior using a 4-parameter sigmoidal (or logistic) curve. These trends were plotted amongst the data points to observe longitudinal trends in data.

### Biological Age Assessment

To estimate biological age from physiological features, we applied principal component analysis (PCA) to three categories of metrics: biomedical (PA mechanics), lung, and right ventricular (RV) measurements. PCA reduces the dimensionality of the dataset by linearly transforming a set of possibly correlated variables into a smaller set of new orthogonal variables, i.e. principal components (PCs), that capture as much variation across samples as possible. The prcomp() function in R was used to conduct PCA with scale. = TRUE, which scales each variable to have unit variance before the analysis. The derived principal components were sorted in decreasing proportion of explained variance. We took the top PCs with a summed proportion of explained variance higher than 60%. Each sample was projected into the subspace spanned by the chosen PCs and the coordinates of all samples on each PC were taken as the estimated biological ages. Clinical traits that formed each PC confirmed the clinical relevance of the chosen PCs. In addition, these PC scores were plotted against chronological age to examine their relationship and assess their validity as biological age indicators. We aggregated multiple observations for samples with repeated measurements by calculating the mean PC1 and PC2 scores across all replicates to yield a single value per sample.

### Single Cell RNA Sequencing (scRNA-seq) and Analyses

We conducted an analysis of viable cells from the primary pulmonary arteries of both young and old mice. Post-euthanasia, we excised the heart, lungs, and pulmonary arteries along with the surrounding tissues. The arteries were mechanically dissected and incubated in a solution of 1 mg/ml collagenase Type I (gibco 17100-017), 1 mg/ml elastase (Worthington, LS002292), dispase II (Sigma-Aldrich, D4693-1G), DNase I (Roche, 101041159001), and hyaluronidase (Sigma, H3506) for 30 minutes. The cells were filtered through a 40 micron mesh, and viable cells were counted using Trypan blue and a Countessa II. Upon achieving cellular dissociation, we utilized the 10x Genomics Chromium platform (3’ v3.1 kit) and a droplet-based microfluidic system, as previously described^7^, to barcode unique mRNA molecules of each cell. Subsequent steps included reverse transcription, cDNA amplification, fragmentation, adaptor ligation, and sample index PCR in accordance with the manufacturer’s instructions. We performed quality control by assessing high sensitivity DNA bioanalyzer traces of cDNA post-barcoding and of the final cDNA library. The final cDNA libraries were sequenced on a HiSeq 4000 Illumina platform in our core facility, targeting 150 million reads per library. Raw sequencing reads were demultiplexed based on sample index adaptors incorporated in the final step of cDNA library preparation. Cutadapt was used to remove any potential adaptor and/or primer contamination. We processed the trimmed reads with the scRNA-seq implementation of STAR (STARsolo), where reads were aligned to the murine reference genome GRCm38 release M22 (GRCm38.p6), collapsed, counted, and summarized into a gene expression matrix. Data analysis and visualization were executed using the Seurat R package26^26^. In detail, we clustered cellular transcriptomes and visualized them in Uniform Manifold Approximation and Projection (UMAP) space to identify cell types (Supplemental Figure S3).

Within each cell type, biological ages were associated with gene expression over the adult lifespan of the mice using a generalized linear mixed effects (GLME) model, which assumes that read counts of a gene across all cells follow a negative binomial distribution with a mean dependent on the biological age and a random intercept grouped by subjects. We included only genes expressed in more than 5% of cells in at least three subjects for the association analysis. Genes that failed to achieve model convergence were excluded. Ultimately, we selected genes with a false discovery rate less than 0.05 as significant biological age correlated genes (BCGs). To elucidate the biological functions of the significant BCGs, we performed pathway enrichment analysis using fGSEA^22^ across four cell types: endothelial cells (ECs), smooth muscle cells (SMCs), fibroblasts (FBs), and myeloid cells.

### Data Availability

Single-cell RNAseq data generated for this study will be available from NCBI Gene Expression Omnibus upon peer-review publication.

## Results

### Organ-specific metrics of age-related changes of the cardiopulmonary system comprise a complex network of structural and functional changes

We found that measures of chronological, age-related, and organ-level changes comprise both linear and non-linear trends (Figure 1). We found that RV contractility decreases over the adult lifespan of mice (Figure 1A top panel). RV adaptation to changes in pulmonary circulation can be heterometric (characterized structurally by RV and RA dilatation) or homeometric (characterized by RV free wall thickness). We found that RV free wall thickness in mice increased later in life, after 24 months (Figure 1A middle panel). Signs of right-sided volume overload occurred earlier in life at ∼20 months (Figure 1A bottom panel). This suggests that decreased contractility of the RV reflects intrinsic aging of the heart, whereas RV remodeling resulting in wall thickening and atrial and ventricular enlargement may be in response to extrinsic factors from other components of the cardiopulmonary system. Individual cardiac measurements for each time point (along with all other physiologic measurements below) are provided in Supplemental Table S1.

**Figure 1.**
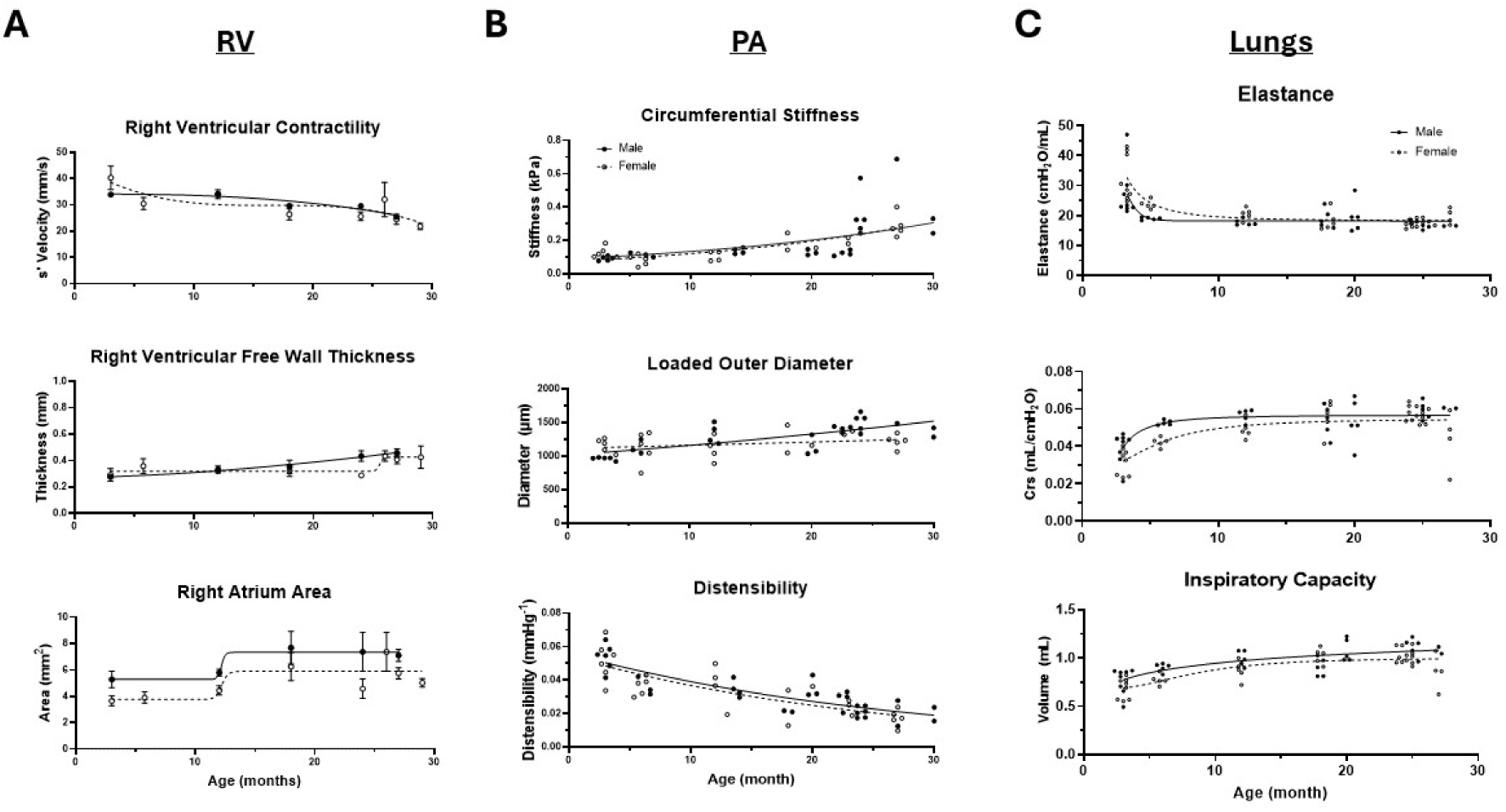

We found that age-related changes in the PA are largely linear in nature. Circumferential stiffness increased as mice increased in age (Figure 1B top panel). There was little change in the thickness of the wall of the PA during the adult life of mice (Supplemental Figure S1), a phenomenon we found to be true previously when comparing young and old mice^7^, and in longitudinal measurements of human PAs^23^. Loaded diameter increased as age increases (Figure 1B middle panel). Distensibility decreased with age (Figure 1B bottom panel), corresponding to a linear increase in PWV (Supplemental Figure S1).

We found that Lung parenchymal elastance decreased rapidly during early adulthood (∼3-5 months) and very gradually declined throughout the lifespan of mice (Figure 1C top panel). Dynamic compliance of the airways increased rapidly during young adulthood and then gradually increased throughout the lifespan of mice (Figure 1C middle panel). The inspiratory capacity of the lungs of mice gradually increased throughout the lifespan of mice (Figure 1C bottom panel).

Trends of RV, PA, and lung mechanical changes were similar with few exceptions. First, lung mechanical measurements tended to be greater in male mice rather than female mice (Figure 1C). We previously found this to be due to allometric differences associated with body mass rather than sex as a biological variable in developing mice^24^. Second, right atrial area tended to be larger in male mice than female mice at all time points (Figure 1A bottom panel). Last, there is a notable increase in RV free wall thickness of 24 month old male mice compared to 24 month old female mice (Figure 1A middle panel).

### Biological aging can be well-characterized by key, organ-related metrics of the RV, PA, and lungs through principal component analyses

To simplify our complex aging data of the cardiopulmonary system, we performed principal component analyses of its major organs, the heart, pulmonary artery, and lungs. Principal component analysis is a mathematical method of transforming complex data to orthogonal linear combinations of the original features to enable data visualization, noise reduction and feature selection. We found that the complex nature of organ-level changes in the cardiopulmonary system can be reduced in complexity to one or two principal components per organ. These changes transform the multiple measurements per organ based on age as a chronological variable to a physiological-based measure of biological aging. Further, this method revealed differences when using sex as a biological variable in select cases. Principal component analysis of the 12 measurements of the RV (Supplemental Table S1) revealed two principal components that characterize aging (Figure 2A). The first component found that changes in RV free wall thickness, RA area, and PA diameter are strong predictors of aging of the RV. The second component suggests that RV contractility strongly associates with aging (TAPSE or tricuspid annular plane systolic excursion; and RVs, which is s’, the tissue doppler measurement of peak velocity of the lateral aspect of the annulus of the tricuspid valve during systole). PCA of the 11 PA measurements (Supplemental Table S1) revealed that mechanical stored energy and stiffness of the arterial wall of the pulmonary artery are associated best with aging of the pulmonary artery. The second component identified, the loaded outer diameter of the pulmonary artery is associated with aging (Figure 2B). Finally, biological aging of the lung was reduced to a single descriptor (Figure 2C), reducing all 8 measurements of the mechanical properties of the lung measured by PFT (Supplemental Table S1) to a single principal component that comprises contributions from static and dynamic compliance (Crs and Cst, respectively), parenchymal elastance (Ers), inspiratory capacity (IC), tissue damping (G), and H (the ratio of the airway resistance to the total respiratory system resistance, where H∼0 is interpreted as the majority of air resistance during forced oscillations results from the airways as seen in healthy lungs, and H∼1 is interpreted as the majority of resistance comes from the lung parenchyma as may be seen in obstructive physiology).

**Figure 2.**
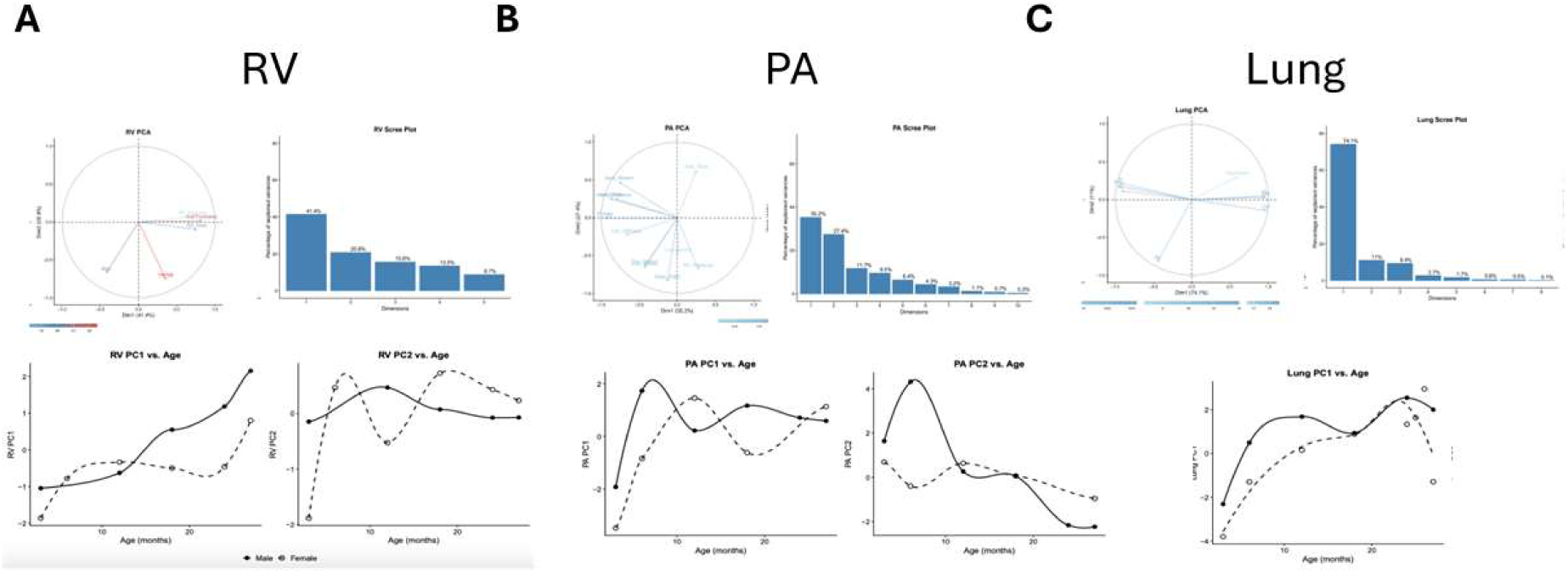

**Figure 3.**
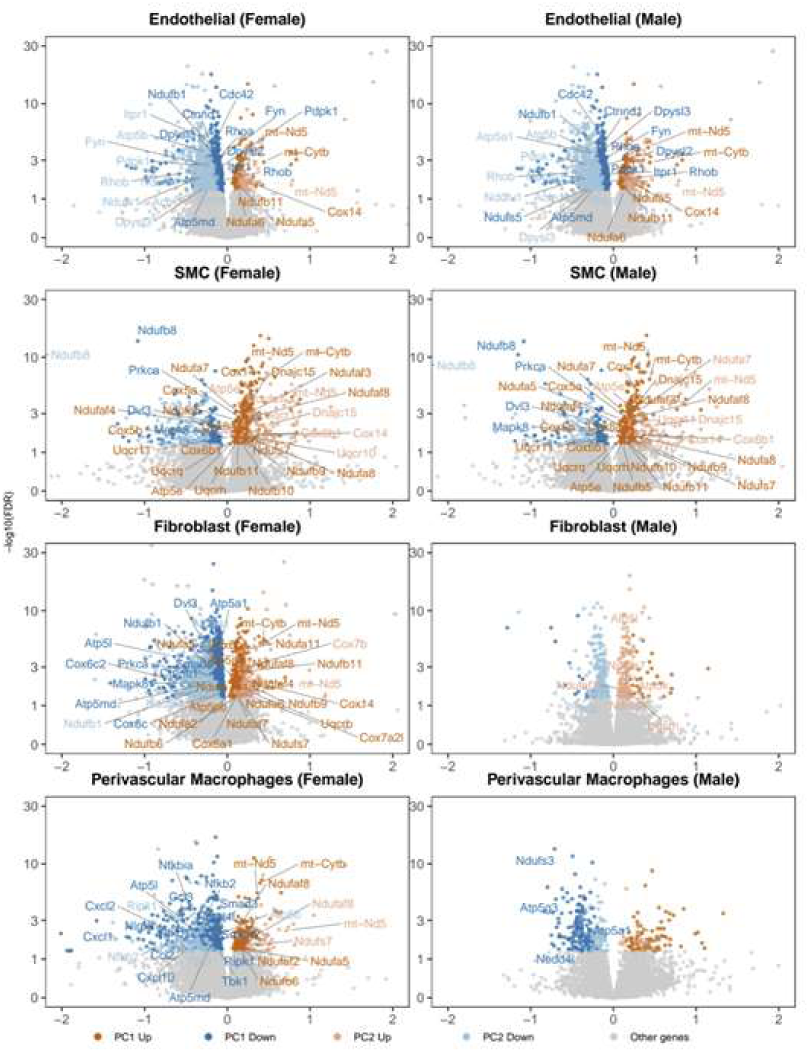

**Figure 4.**
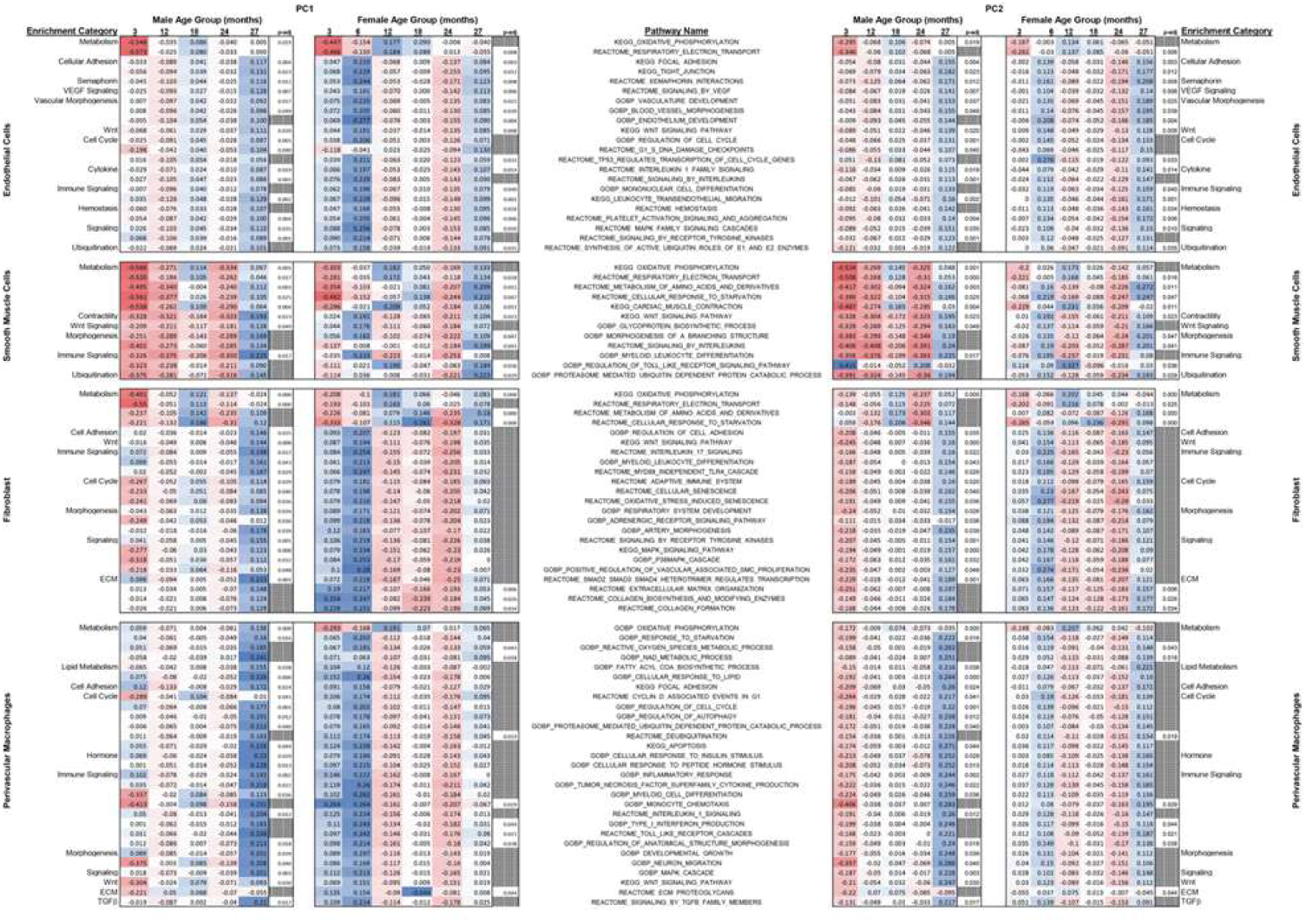

### Biological measures of aging rather than chronological aging reveal transcriptomic cell-specific molecular mechanisms of aging of the cardiopulmonary system

We isolated 132,997 cells from 19 samples of PAs from mice ranging from 3 months to 28 months, 9 female and 10 male (Supplemental Table S1). 7 of these samples we reported previously^7^.

We embedded these cells in UMAP and defined our cells with gene markers (Supplemental Figure S3). Attempts to identify gene expression as a function of age as a chronological variable yielded few genes of unclear physiological significance to aging (Supplemental Figure S2). Identifying biological aging associated genes (BAAGs) using biological aging identified from our PCAs yielded 13,636 significant genes that are physiologically meaningful.

### Aging associates with expression of genes associated with oxidative phosphorylation throughout cells that reside in the arterial wall and cell-specific decreases in mechanotransduction, Wnt signaling, and TGFb/NFkB pathways

Amongst EC, SMC, Fib, and Perivascular Mac, there is significant increase in expression of genes associated with oxidative phosphorylation, including mt-Nd5, mt-Cytb, Cox, and Nduf, Uqcr, Atp5 subunits, Danjc15, and Park7. In EC, there is decreased expression of genes associated with Focal adhesion (Rhoa, Cdc42, Vcl, Actn1), VEGF signaling (Fyn, Pdpk1, Itpr1), and semaphorin guidance (Dpysl2/3, Rhob). Reduction in these processes indicate reduced mechanotransduction signaling, which leads to loss of mechano-adaptive programming^25,26^. Uncoupling mitochondrial respiration from shear stress sensing has been shown to elevate superoxide production, reducing NO bioavailability, and increases pulmonary vascular resistance^27,28^. There are also decreased expression of genes associated with reduced EC barrier (Ctnnd1, Ctnnb1, Tjp2, Vcl). In SMC, there is significant decrease in genes associated with Wnt signaling (Prkca, Dvl3, Mapk8, Ctnnb1, Lrp5/6, Smad4, Fgfr2, Esr1, and Yap1).

In adventitial fibroblasts, there was a significant decrease in expression of genes associated with Wnt signaling (Dvl3, Gsk3b, Tle1, Tbl1x/Ppp2ca), MAPK signaling (Egfr, Shc1, Dusp16/Dusp7, Calm1, Bcl2l1). and cell cycling and senescence (Mapk1, Fox, Ccnd2, Rptor, Cdc42, Brd4, and Ywhab/z/q). Genes associated with matrix organization include collagens (Col1a1, Col6a1, Col12a1, Col15a1) and matrix remodeling genes including Adamts2 (involved in collagen maturation), Plod1 (involved cross-linking of collagen), and Bmp1.

In perivascular macrophages, coordinated downregulation of innate immune and stress-response programs suggests a shift toward dysfunctional vascular wall remodeling. Key nodes in TLR/NF-κB signaling (Nfkbia, Nfkb1/2, IRAK1/2, Ripk1/2, Tbk1) are decreased, consistent with reduced lipopolysaccharide-mediated signaling and blunted inflammatory cytokine/chemokine output (Ccl2, Ccl3, Cxcl1/2/10, TNF, IL1B), alongside reduced myeloid/lymphocyte differentiation pathways. In parallel, lower expression of genes linked to lipid oxidation and inflammasome/uptake signaling (Nlrp3, Ldlr, Prkca, Tgfb1) and diminished TGFβ-family signaling capacity (Smad3, Smad4, Tgfb1, Nedd4l, Smurf1/2) would be expected to weaken macrophage-to–smooth muscle cell (SMC) and endothelial cell (EC) crosstalk that normally stabilizes SMC contractile tone and maintains EC barrier homeostasis. Functionally, reduced TNF/IL-1β and chemokines can limit compensatory vasodilatory signaling and impair macrophage–EC communication needed under hypoxic or altered hemodynamic stress (notably via decreases in Cxcl10/Cxcr4 and Ccl2), while retained oxidative phosphorylation may promote mitochondrial ROS that indirectly activates endothelial NF-κB and endothelial dysfunction. Finally, because Nlrp3 contributes to physiologic IL-18/IL-1β-dependent regulation of vascular tone, reduced Nlrp3 signaling could remove a brake on maladaptive SMC growth, potentially permitting unchecked SMC hypertrophy and contributing to age-associated vascular wall thickening and remodeling.

Overall, these transcriptomic changes are similar to those that we observed when analyzing transcriptomic changes using age as a categorical variable, young versus old^7^. However, treating aging as a biological variable rather than a categorical or chronological variable enabled us to observe transcriptomic changes more closely aligned with the process and biology of aging rather than old age.

## Discussion

We hypothesized that cardiopulmonary aging is nonlinear across the lifespan of mice, where cardiopulmonary aging occurs in phases rather than uniform trajectories. We further hypothesized that the PA is an active participant in age-related cardiopulmonary decline resulting in a positive feedback loop whereby PA stiffening increases RV afterload and pulmonary hemodynamics contributing to a decline in both RV and lung function. We found that aging of the RV, PA, and lungs are tightly coupled, such that cardiopulmonary decline emerges from interdependent, linear and non-linear age-related changes rather than a simple function of chronological time. Organ-specific metrics capturing structural and functional remodeling across the RV, PA, and lungs form an integrated network, reflecting coordinated changes in ventricular adaptation, pulmonary vascular mechanics, and parenchymal function. Leveraging these organ-resolved metrics, biological aging can be quantitatively reduced to summarize shared and tissue-specific aging variance, in other words, physiologic-based biological aging of the cardiopulmonary system. Interpreting transcriptomic remodeling as a function of biological—rather than chronological—age reveals cell-type–specific mechanisms that underpin cardiopulmonary aging, enabling causal hypotheses that link molecular programs to emergent organ-level dysfunction.

### Physiologic-based biological aging of the cardiopulmonary system

Age-related changes in the cardiopulmonary system are neither uniformly linear nor non-linear. The pulmonary artery Exhibited largely linear stiffening and decreased distensibility across the adult lifespan this is consistent with the progressive cumulative nature of structural material extracellular matrix remodeling in the aorta.^29-33^ Similarly, the RV exhibited largely linear decline in contractility as age increased. These appear to be intrinsic age-related properties of the PA and RV. In contrast, the lung and RV exhibited non-linear patterns of age-related change with early-adulthood change in lung parenchyma mechanism followed by a plateau then gradual decline. These changes likely reflect post-developmental alveolar maturation followed by gradual, progressive loss of alveolar units and their associated microvasculature^34-36^. Non-linear age-related RV changes are consistent with compensatory remodeling to increased afterload appear to accelerate only late in life. Therefore, the RV displays a two-stage model of aging: an early, intrinsic stage driven by linear, progressive loss of cardiomyocyte contractility; and a later, extrinsic stage in which RV remodeling (RV free wall thickening and right atrial enlargement) is drive by increasing afterload from a progressive stiffer pulmonary vasculature exacerbated by rarefaction of the aging lung due to alveolar remodeling and loss of pulmonary microcirculation. These are consistent with models of RV-PA coupling^37-39^ and findings that RV hypertrophy and dilation are load-sensitive responses that occur slower than intrinsic contractility dysfunction^40^. The implications of these findings are that much of the structural setting of lung mechanical reserve occurs early in life and that early RV contractile decline may be a sensitive and potentially reversible marker of biological aging. Interventions aimed at preserving lung and RV function may need to be initiated far earlier than currently appreciated.

The sex differences observed in RV free wall thickness at 24 months — with males showing greater hypertrophy than females — are noteworthy and may reflect sex-specific differences in RV adaptive capacity or in the magnitude of pulmonary vascular afterload. These findings are consistent with prior reports of sex-specific RV remodeling in response to pressure overload^41,42^, and suggest that biological aging of the RV may be sexually dimorphic.

This divergence of intrinsic and extrinsic aging of individual components suggests that a single chronological variable may not be sufficient to adequately represent aging of the entire cardiopulmonary system. This is consistent with evidence that different organs age at different rates in biological aging studies in humans^43-45^. We extend this concept to the cardiopulmonary system in a longitudinal model providing mechanistic insights that human populations studies cannot easily provide. The anatomical and physiological linker of the RV and lungs is the PA. Our finding that PA stiffening (and decreased distensibility) is largely linear with age—in contrast to patters of the RV and lung—positions the PA as a consistent and measure indicator of cardiopulmonary biological age. PA stiffening that may be more amenable to continuous monitoring and extrapolation, suggesting that PA mechanical properties may serve a a reliable longitudinal biomarker as we previously proposed^46^. We found that the PA wall thickness did not change significantly with age, consistent with our prior work using mice and human studies^7,23^, suggesting that stiffening is drive primarily by material property changes rather than structural properties such as excessive deposition of collagen and significant wall thickening. This is important because material stiffening can occur without overt structural changes visible on conventional imaging, meaning that standard anatomical assessments of the pulmonary artery may significantly underestimate the degree of vascular aging present. Yet, wall stiffening results in increased PWV, which increased linearly with our age in our data, is an established clinical measure of arterial stiffness in the systemic circulation and independently predictive of cardiovascular outcomes^47^.

### Transmural metabolic-inflammatory uncoupling associates with suppressed adaptive/regulatory signaling (mechanotransduction, WNT, TGFβ/NFκB), promoting arterial wall stiffening and decreased NO-mediated vasodilation

Perhaps the most methodologically significant finding of this study is that using PCA-derived biological aging as the continuous independent variable to correlate gene expression with biological cage identified physiologically meaningful transcriptomic changes, which we have called biological age correlated genes (BCGs). This finding has broad methodological implications for the field of vascular aging and single-cell transcriptomics. The likely explanation for this is that biological aging integrated multiple functional dimension of organs and separate components of the cardiopulmonary system, thereby reducing the non-biological noise contributed by individual variation in chronological age, similar to the use of epigenetic clocks to replace chronological age in human aging studies^48,49^.

Using biological aging, we found that age-related EC metabolic dysregulation associates with impaired mechanotransduction. There is a paradoxical enrichment of oxidative phosphorylation genes in conjunction with decreased leukocyte transendothelial migration, VEGF signaling, and blood vessel morphogenesis pathways consistent across both sexes. The concurrent oxidative phosphorylation increases with decreased mechanosensory adhesion and permeability genes mirrors to metabolic shifts as seen in the Warburg Effect as seen in EC’s in PAH as well as the eNOS-RhoA-VEGFR2 signaling cascades. SMC oxidative phosphorylation increase couple with WNT suppression reflects a phenotypic shift from contractile to synthetic phenotype that associates with medial stiffening. Wnt-related SMC phenotypic switching is well-associated with arterial wall stiffening. Oxidative phosphorylation increase may be viewed as a reaction to protect contractile phenotype, which would reduce wall stress. Park7 (DJ-1) is a well-known gene associated with mitochondrial protection in vascular SMC’s under oxidative stress; whereas, synthetic phenotype is a maladaptive response. Perivascular macrophages mediate immune activation that impairs vascular tone through TGFβ/NFκB-mediates suppression of vasodilatory signing. Metabolic shifts in perivascular macrophages associates with Nlrp3/TGFβ/SMAD paracrine signaling with SMC’s^50-52^. As a result changes in oxidative phosphorylation in perivascular macrophages promote anti-inflammatory but pro-fibrotic pathways. While metabolic alterations have been strongly linked to age-related senescence, our findings provide granular, cell-specific mechanisms of age-related changes within the arterial walls that lead to pulmonary artery stiffening. Thus, the implications of our findings are these cell-specific molecular pathways are potential targets to prevent or delay age-related stiffening of the PA, which may delay age-related cardiopulmonary functional decline.

### The proximal pulmonary artery is an active contributor to Cardiopulmonary Aging

Taken together, the structural, functional, and transcriptomic findings of this study support the hypothesis that the PA is an active participant in — not merely a reflection of — cardiopulmonary aging. PA stiffening increases RV afterload promotes pulsatile pressure transmission to the pulmonary microcirculation, and impairs RV-vascular coupling. These hemodynamic consequences feed back onto RV structure and function and, through altered shear stress and pressure loading, likely further accelerate PA remodeling. This creates a positive feedback loop that must be broken to delay or prevent aging.

Our transcriptomic data add molecular insights to this feedback loop. Loss of EC mechanotransduction impairs the PA’s ability to adapt to altered hemodynamics, accelerating dysfunction. Loss of SMC Wnt signaling reduces contractile reserve, further stiffening the vessel. Macrophage signaling erosion impairs the vascular wall’s capacity for organized repair. Each of these cell-type-specific changes reduces the arterial wall’s ability to store elastic energy and buffer pulse wave velocity and pulsatility from penetrating deep into the pulmonary vascular bed. Thus stiffness is a homeostatic setpoint that is gradually altered during aging, making it progressively less capable of responding to the very hemodynamic perturbations that its own stiffening creates.

### Limitations

We derived this physiological and transcriptomic data from mice and therefore require direct validation in human tissue. While this research design is quasi-longitudinal, the terminal nature of experiments makes is cross-sectional in nature. Therefore, we cannot infer causality from association between biological aging, physiological changes, and molecular-cellular changes that we observed. However, these insights provide targets for investigation in longitudinal cohort for future study. Our focus on the proximal pulmonary artery mechanics identify the need to investigate distal vascular changes in separate studies. Our PCA-derived method for measuring biological aging were internally derived and require validation against independent measures of biological age such as epigenetic clock and plasma-proteomics based clocks. Further, we previously found that microstructural remodeling was a primary driver of PA stiffening^15,53^. Therefore, it is important to evaluate the potential benefit of incorporating microstructure of the proximal pulmonary artery into biomechanical age-related changes to determine if this improves measuring biological age. Last, the identification of altered metabolism, signaling erosion, and macrophage dysfunction as candidate mechanisms of PA aging provides a roadmap for targeted interventional studies — using, for example, anti-aging metabolic altering agents such as metformin and rapalogs, senolytic agents, or myeloid replacement — to test whether reversing these molecular changes can restore more youthful biomechanical properties.

### Conclusion

The cardiopulmonary system ages as an integrated, inter-dependent unit, in which the mechanical properties of the pulmonary vasculature, the adaptive capacity of the RV, and the structural integrity of the lung parenchyma are continuously coupled through hemodynamic, mechanical, and cellular-molecular crosstalk. Chronological age captures only a fraction of these biological phenomena. By grounding the study of cardiopulmonary aging in physiologic, organ-specific measures of biological aging, this work demonstrates that the molecular mechanisms of aging are more coherently revealed, more biologically interpretable, and more likely to yield actionable therapeutic targets than analyses anchored to calendar time alone. Future work should aim to translate these findings to human tissue and populations, validate biological aging scores as clinical biomarkers, and test whether identified molecular pathways and cell-directed therapies represent reversible targets for cardiopulmonary rejuvenation.

---

**Supplemental Table S1.**
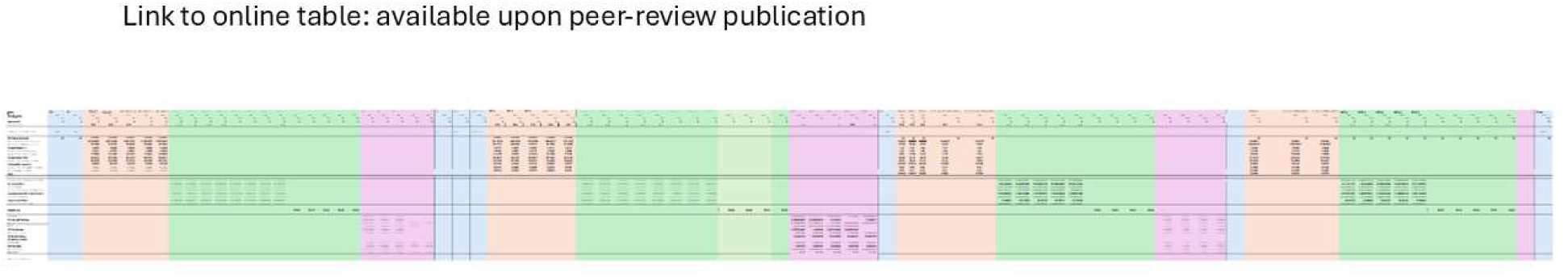

**Supplemental Figure S1:**
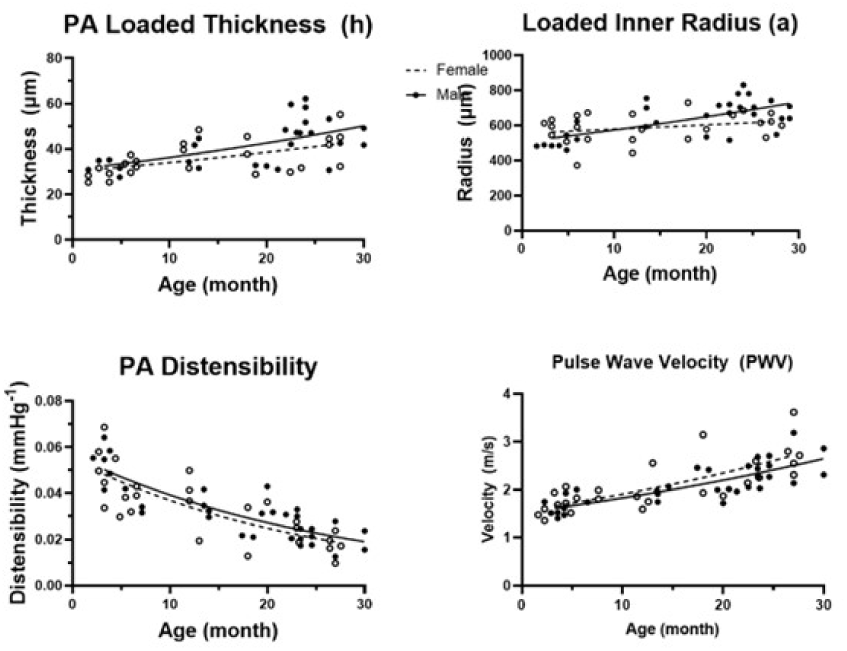
additional physiological data.

**Supplemental Figure S2:**
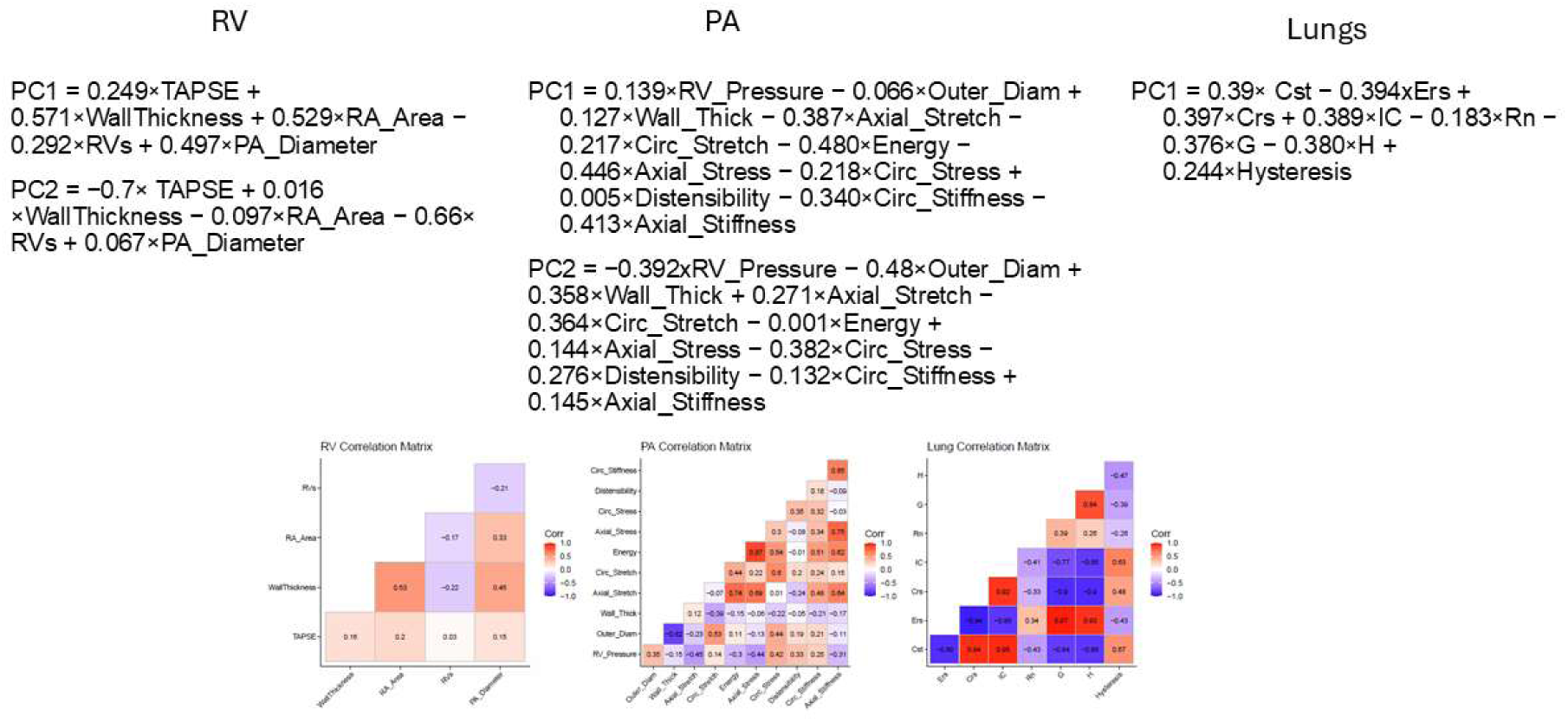
PCA supplemental data.

**Supplemental Figure S3.**
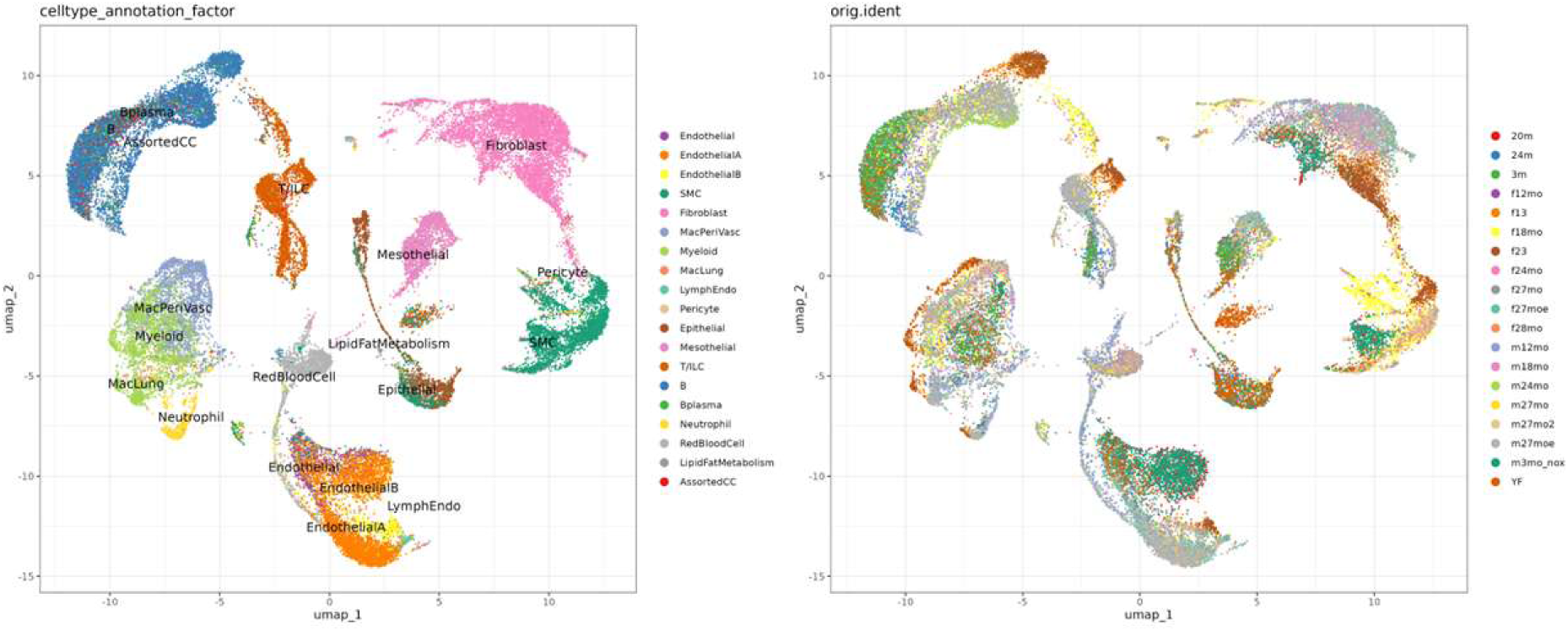
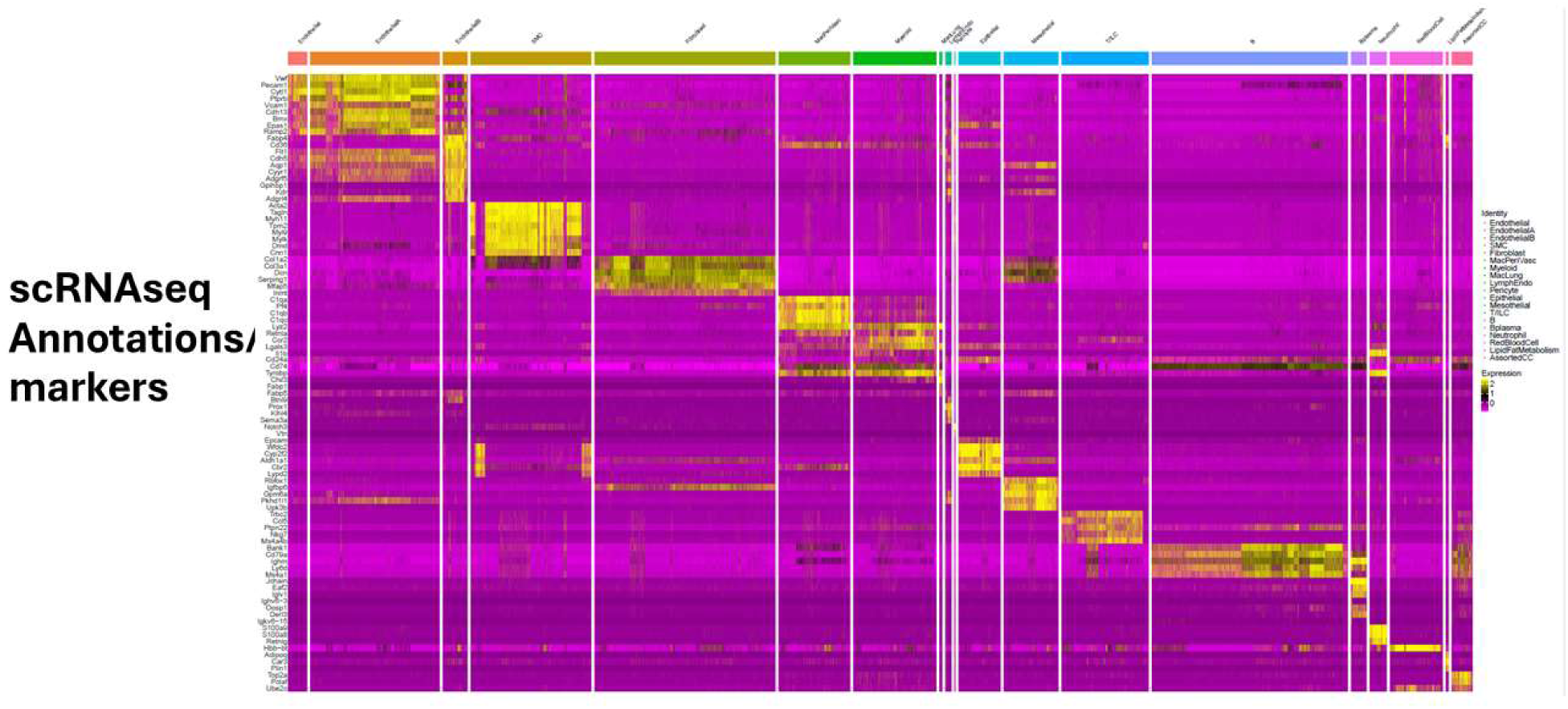

**Supplemental Table S2.**
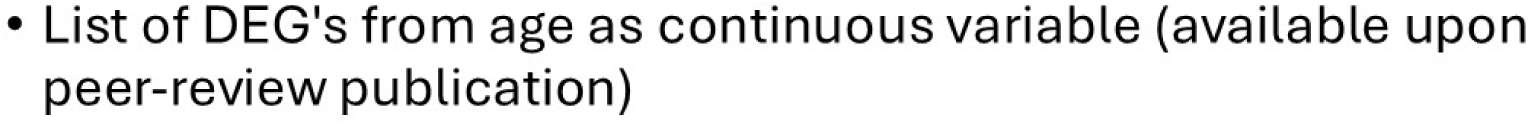

**Supplemental Table S3.**
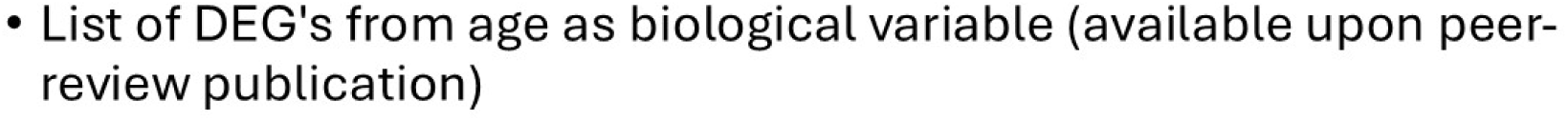

**Supplemental Figure S4.**
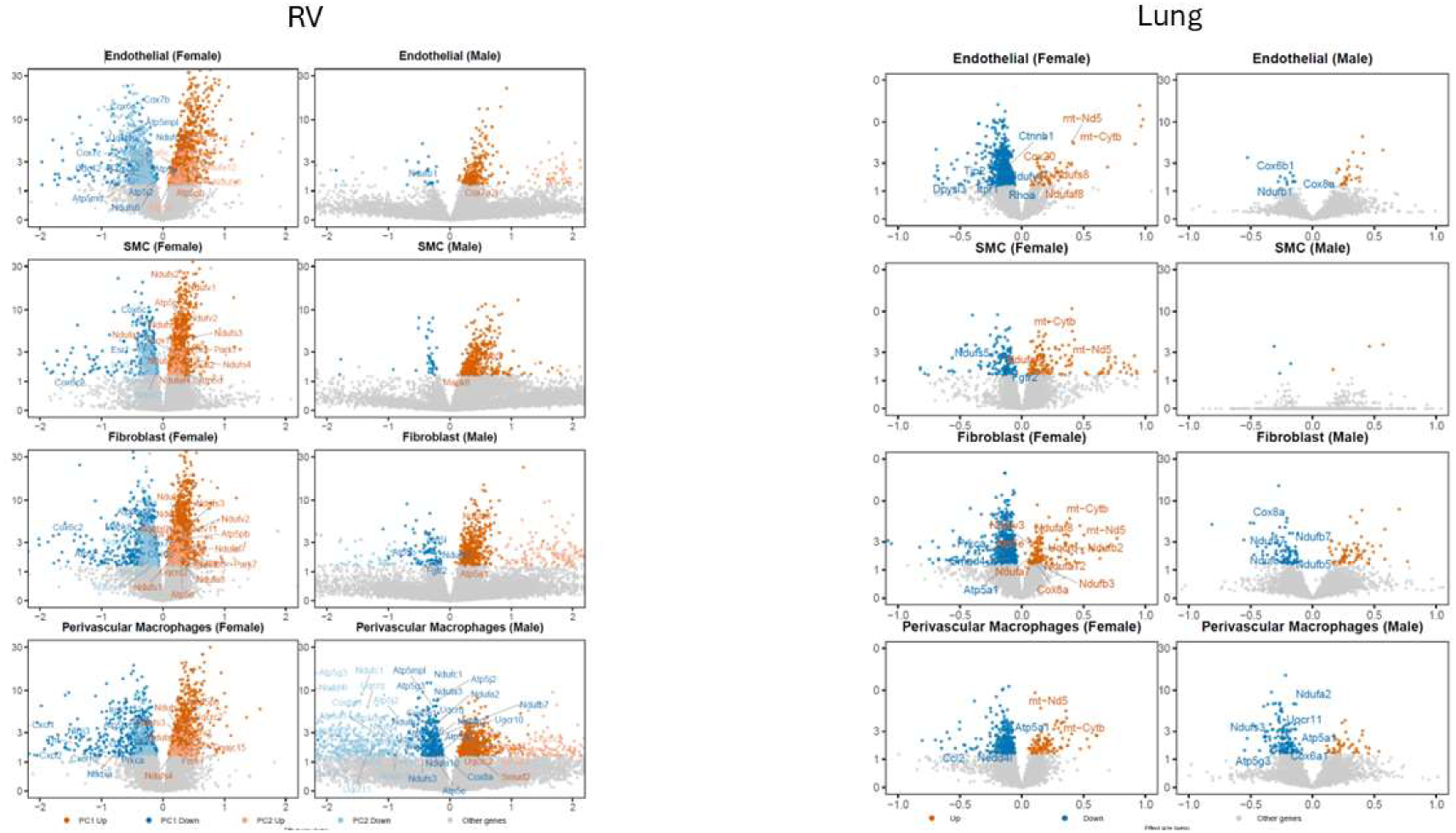

## Notes

Disclosures: This work was supported by the VA VISN1 Fred Wright CDA1, National Institute on Aging R03AG074063, and EPM is a Pepper Scholar of the Yale Claude D. Pepper Older Americans Independent Center supported by NIA P30AG021342. EPM is a consultant for Biomedical Consultants, PLLC. ABR is supported by NSF grant 2436623.

### Competing Interest Statement

The authors have declared no competing interest.

## References

1. Lakatta EG, Levy D. Arterial and cardiac aging: major shareholders in cardiovascular disease enterprises: Part I: aging arteries: a “set up” for vascular disease. Circulation. 2003;107(1):139–146.

2. Lakatta EG, Levy D. Arterial and cardiac aging: major shareholders in cardiovascular disease enterprises: Part II: the aging heart in health: links to heart disease. Circulation. 2003;107(2):346–354.

3. Boutouyrie P, Bussy C, Lacolley P, Girerd X, Laloux B, Laurent S. Association Between Local Pulse Pressure, Mean Blood Pressure, and Large-Artery Remodeling. Circulation. 1999;100(13):1387–1393.

4. Boutouyrie P, Tropeano AI, Asmar R, et al. Aortic stiffness is an independent predictor of primary coronary events in hypertensive patients: a longitudinal study. Hypertension. 2002;39(1):10–15.

5. Laurent S, Boutouyrie P, Asmar R, et al. Aortic stiffness is an independent predictor of all-cause and cardiovascular mortality in hypertensive patients. Hypertension. 2001;37(5):1236–1241.

6. Horvath S, Raj K. DNA methylation-based biomarkers and the epigenetic clock theory of ageing. Nature reviews genetics. 2018;19(6):371–384.

7. De Man R, Cai Z, Doddaballapur P, et al. Proximal Pulmonary Artery Stiffening as a Biomarker of Cardiopulmonary Aging. Aging Cell. 2026;in press.

8. Beckers J, Wurst W, de Angelis MH. Towards better mouse models: enhanced genotypes, systemic phenotyping and envirotype modelling. Nature Reviews Genetics. 2009;10(6):371–380.

9. Wahlsten D, Metten P, Phillips TJ, et al. Different data from different labs: Lessons from studies of gene–environment interaction. Journal of Neurobiology. 2003;54(1):283–311.

10. Ramachandra AB, Humphrey JD. Biomechanical characterization of murine pulmonary arteries. Journal of biomechanics. 2019;84:18–26.

11. Hanks JH, Wallace RE. Relation of oxygen and temperature in the preservation of tissues by refrigeration. Proceedings of the Society for Experimental Biology and Medicine. 1949;71(2):196–200.

12. Gleason R, Humphrey J. A mixture model of arterial growth and remodeling in hypertension: altered muscle tone and tissue turnover. Journal of vascular research. 2004;41(4):352–363.

13. Ferruzzi J, Bersi M, Humphrey J. Biomechanical phenotyping of central arteries in health and disease: advantages of and methods for murine models. Annals of biomedical engineering. 2013;41(7):1311–1330.

14. Humphrey JD. Cardiovascular solid mechanics: cells, tissues, and organs: Springer Science & Business Media; 2013.

15. Manning EP, Ramachandra AB, Schupp JC, et al. Mechanisms of Hypoxia-Induced Pulmonary Arterial Stiffening in Mice Revealed by a Functional Genetics Assay of Structural, Functional, and Transcriptomic Data. Frontiers in Physiology. 2021;12(1510).

16. Bramwell JC, Hill AV. The velocity of pulse wave in man. Proceedings of the Royal Society of London. Series B, Containing Papers of a Biological Character. 1922;93(652):298–306.

17. Sanz J, Prat-Gonzalez S, Macaluso F, Fuster V, Garcia M. 155 Quantification of pulse wave velocity in the pulmonary artery in patients with pulmonary hypertension. presented at: Journal of Cardiovascular Magnetic Resonance2008.

18. Gupta A, Sharifov OF, Lloyd SG, et al. Novel noninvasive assessment of pulmonary arterial stiffness using velocity transfer function. Journal of the American Heart Association. 2018;7(18):e009459.

19. Peng H-H, Chung H-W, Yu H-Y, Tseng W-YI. Estimation of pulse wave velocity in main pulmonary artery with phase contrast MRI: Preliminary investigation. Journal of Magnetic Resonance Imaging. 2006;24(6):1303–1310.

20. Lindsey ML, Kassiri Z, Virag JA, de Castro Brás LE, Scherrer-Crosbie M. Guidelines for measuring cardiac physiology in mice. American Journal of Physiology-Heart and Circulatory Physiology. 2018;314(4):H733–H752.

21. Kohut A, Patel N, Singh H. Comprehensive Echocardiographic Assessment of the Right Ventricle in Murine Models. J Cardiovasc Ultrasound. 2016;24(3):229–238.

22. Korotkevich G, Sukhov V, Budin N, Shpak B, Artyomov MN, Sergushichev A. Fast gene set enrichment analysis. bioRxiv. 2021:060012.

23. Manning EP, Mishall P, Ramachandra AB, et al. Stiffening of the human proximal pulmonary artery with increasing age. Physiol Rep. 2024;12(12):e16090.

24. Bärnthaler T, Ramachandra AB, Ebanks S, et al. Developmental changes in lung function of mice are independent of sex as a biological variable. Am J Physiol Lung Cell Mol Physiol. 2024.

25. Romani P, Valcarcel-Jimenez L, Frezza C, Dupont S. Crosstalk between mechanotransduction and metabolism. Nature Reviews Molecular Cell Biology. 2021;22(1):22–38.

26. Gargalionis AN, Papavassiliou KA, Papavassiliou AG. Mechanotransduction Circuits in Human Pathobiology. Int J Mol Sci. 2024;25(7).

27. Hsieh HJ, Liu CA, Huang B, Tseng AH, Wang DL. Shear-induced endothelial mechanotransduction: the interplay between reactive oxygen species (ROS) and nitric oxide (NO) and the pathophysiological implications. J Biomed Sci. 2014;21(1):3.

28. Jones CI, 3rd, Han Z, Presley T, et al. Endothelial cell respiration is affected by the oxygen tension during shear exposure: role of mitochondrial peroxynitrite. Am J Physiol Cell Physiol. 2008;295(1):C180–191.

29. Humphrey JD, Harrison DG, Figueroa CA, Lacolley P, Laurent S. Central Artery Stiffness in Hypertension and Aging: A Problem With Cause and Consequence. Circ Res. 2016;118(3):379–381.

30. Laurent S, Cockcroft J, Van Bortel L, et al. Expert consensus document on arterial stiffness: methodological issues and clinical applications. European Heart Journal. 2006;27(21):2588–2605.

31. Chirinos JA, Segers P, Hughes T, Townsend R. Large-artery stiffness in health and disease: JACC state-of-the-art review. Journal of the American College of Cardiology. 2019;74(9):1237–1263.

32. Townsend RR, Rosendorff C, Nichols WW, et al. American Society of Hypertension position paper: central blood pressure waveforms in health and disease. Journal of the American Society of Hypertension. 2016;10(1):22–33.

33. Herzog MJ, Müller P, Lechner K, et al. Arterial stiffness and vascular aging: Mechanisms, prevention, and therapy. Signal transduction and targeted therapy. 2025;10(1):282.

34. Janssens J-P, Pache J-C, Nicod L. Physiological changes in respiratory function associated with ageing. European Respiratory Journal. 1999;13(1):197–205.

35. Schulte H, Mühlfeld C, Brandenberger C. Age-related structural and functional changes in the mouse lung. Frontiers in physiology. 2019;10:1466.

36. Brandenberger C, Mühlfeld C. Mechanisms of lung aging. Cell and tissue research. 2017;367(3):469–480.

37. Vonk Noordegraaf A, Westerhof BE, Westerhof N. The relationship between the right ventricle and its load in pulmonary hypertension. Journal of the American College of Cardiology. 2017;69(2):236–243.

38. Vonk-Noordegraaf A, Haddad F, Chin KM, et al. Right heart adaptation to pulmonary arterial hypertension: physiology and pathobiology. Journal of the American College of Cardiology. 2013;62(25 Supplement):D22-D33.

39. Thenappan T, Ormiston ML, Ryan JJ, Archer SL. Pulmonary arterial hypertension: pathogenesis and clinical management. Bmj. 2018;360.

40. Rain S, Handoko ML, Trip P, et al. Right ventricular diastolic impairment in patients with pulmonary arterial hypertension. Circulation. 2013;128(18):2016–2025.

41. Jacobs W, van de Veerdonk MC, Trip P, et al. The Right Ventricle Explains Sex Differences in Survival in Idiopathic Pulmonary Arterial Hypertension. CHEST. 2014;145(6):1230–1236.

42. Frump AL, Albrecht M, Yakubov B, et al. 17â-Estradiol and estrogen receptor á protect right ventricular function in pulmonary hypertension via BMPR2 and apelin. The Journal of clinical investigation. 2021;131(6).

43. Nie C, Li Y, Li R, et al. Distinct biological ages of organs and systems identified from a multi-omics study. Cell reports. 2022;38(10).

44. Li Y, Xu X, Zheng Y, et al. Synergistic and heterogeneous aging using composite phenotypes and multiple organ systems aging clocks. Communications Medicine. 2025;5(1):506.

45. Oh HS-H, Rutledge J, Nachun D, et al. Organ aging signatures in the plasma proteome track health and disease. Nature. 2023;624(7990):164–172.

46. De Man R, Cai Z, Doddaballapur P, et al. Proximal Pulmonary Artery Stiffening as a Biomarker of Cardiopulmonary Aging. Aging Cell. 2026;25(2):e70383.

47. Vlachopoulos C, Aznaouridis K, Stefanadis C. Prediction of cardiovascular events and all-cause mortality with arterial stiffness: a systematic review and meta-analysis. Journal of the American College of Cardiology. 2010;55(13):1318–1327.

48. Horvath S. DNA methylation age of human tissues and cell types. Genome biology. 2013;14(10):3156.

49. Lu AT, Quach A, Wilson JG, et al. DNA methylation GrimAge strongly predicts lifespan and healthspan. Aging (albany NY). 2019;11(2):303.

50. Olona A, Leishman S, Anand PK. The NLRP3 inflammasome: regulation by metabolic signals. Trends in Immunology. 2022;43(12):978–989.

51. Wang J, Yuan R, Zhang S, Xu Z, Song L. Metabolic Reprogramming Intermediates of Glucose Regulate Macrophage Polarization: An Important Direction for Ameliorating Pulmonary Vascular Remodeling. Journal of Inflammation Research. 2025;18:16045–16062.

52. Massagué J, Sheppard D. TGF-&#x3b2; signaling in health and disease. Cell. 2023;186(19):4007–4037.

53. De Man R, Cai Z, Doddaballapur P, et al. Proximal Pulmonary Artery Stiffening as a Biomarker of Cardiopulmonary Aging. Aging Cell. 2026;25(2):e70383.

